# Genetic Variability affects the Skeletal Response to Unloading

**DOI:** 10.1101/2020.06.26.174326

**Authors:** Michael A. Friedman, Abdullah Abood, Bhavya Senwar, Yue Zhang, Camilla Reina Maroni, Virginia L. Ferguson, Charles R. Farber, Henry J. Donahue

## Abstract

Mechanical unloading decreases bone volume and strength. In humans and mice, bone mineral density is highly heritable, and in mice the response to changes in loading varies with genetic background. Thus, genetic variability may affect the response of bone to unloading. As a first step to identify genes involved in bone response to unloading, we evaluated the effects of unloading in eight inbred mouse strains: C57BL/6J, PWK/PhJ, WSB/EiJ, A/J, 129S1/SvImJ, NOD/ShiLtJ, NZO/HlLtJ, and CAST/EiJ. C57BL/6J and NOD/ShiLtJ mice had the greatest unloading induced loss of diaphyseal cortical bone volume and strength. NZO/HlLtJ mice had the greatest metaphyseal trabecular bone loss, and C57BL/6J, WSB/EiJ, NOD/ShiLtJ, and CAST/EiJ mice had the greatest epiphyseal trabecular loss. Bone loss in the epiphyses displayed the highest heritability. With immobilization, mineral:matrix was reduced, and carbonate:phosphate and crystallinity were increased. A/J mice displayed the greatest unloading induced loss of mineral:matrix. Changes in gene expression in response to unloading were greatest in NOD/ShiLtJ and CAST/EiJ mice. The most upregulated genes in response to unloading were associated with increased collagen synthesis and extracellular matrix formation. Our results demonstrate a strong differential response to unloading as a function of strain. Diversity outbred (DO) mice are a high-resolution mapping population derived from these eight inbred founder strains. These results suggest DO mice will be highly suited for examining the genetic basis of the skeletal response to unloading.

## Introduction

Unloading of limbs due to prolonged bedrest, immobilization, or exposure to microgravity, decreases bone volume and strength, making bone more susceptible to fracture.^1^ However, the underlying mechanisms that lead to bone loss from unloading remain largely unknown. In both humans and mice, bone mineral density is highly heritable. ^2-5^ Additionally, in mice the response to changes in mechanical loading varies as a function of genetic background.^3,6-11^ Indeed, Judex *et al*. identified six quantitative trait loci (QTLs) for unloading-induced loss of BV/TV in the F2 offspring of a double cross between BALB/cByJ22 and C3H/HeJ23 mice.^11^ These QTL accounted for 21% of the variability in loss of BV/TV in response to unloading in the F2 mice. In a subsequent study, Judex *et al*. identified five QTLs for unloading-induced loss in cortical bone area in the F2 of the same cross.^12^ These QTLs accounted for 10% of the variability in cortical bone loss in response to unloading. These data strongly suggest genetic variability affects the response of bone to unloading. However, identifying which specific genes contribute to the variability in response to unloading is challenging as each QTL contains several hundred candidate genes, and low-resolution mapping approaches have made it difficult to pinpoint causal genes.

Diversity outbred (DO) mice are a recently developed high-resolution mapping population that enables gene discovery for traits such as the response of bone to unloading.^13^ DO mice are derived from eight founder strains and have been outbred for at least 10 generations. DO mice can be used to investigate the genetics of a wide range of complex diseases and may be used to investigtate the response of bone to unloading. Hindlimb suspension (HLS) is the most widely used and effective model for inducing bone loss from unloading.^14-17^ HLS results in decreases in cortical and trabecular bone volume as well as loss of bone structural and tissue strength in as little as three weeks. However, when testing effects of unloading, HLS or other unloading models require a control group of mice that are not unloaded. HLS is not suitable for studying DO mice since the DO mice are not all genetically identical. The single limb immobilization model can potentially overcome this problem by only unloading one limb and using the contralateral limb from the same animal as a control. This model has been used extensively to show decreases in muscle mass and protein synthesis and similar increases in muscle atrophy gene expression, relative to that of HLS after one week.^14,18,19^ We have recently began using single limb immobilization for bone loss and found this model decreases femoral bone volume and bone strength after three weeks.^20^

In anticipation of future studies on DO mice, we employed the single limb immobilization model, rather than HLS, to induce bone loss. Here, we hypothesized that genetic variation would impact the response of bone to unloading after three weeks of single limb immobilization. To test this hypothesis, we evaluated the effects of single limb immobilization in the eight DO mouse founder strains: C57BL/6J, PWK/PhJ, WSB/EiJ, A/J, 129S1/SvImJ, NOD/ShiLtJ, NZO/HlLtJ, and CAST/EiJ. Our results demonstrate a strong differential response as a function of strain and suggest that the DO mice will be highly suited to dissect the genetic basis of the skeletal response to unloading.

## Results

### Effects of Immobilization on Cortical and Trabecular Bone Geometry were Mouse Strain-Dependent

In cortical bone of the mid-diaphysis of the femur, there was a main effect of mouse strain on Ct.Ar, Tt.Ar, Ct.Ar/Tt.Ar, and Ct.Th (p<0.0001), and there was a main effect of immobilization on Ct.Ar/Tt.Ar and Tt.Ar (p<0.05, Figure 1). Among the mouse strains, Ct.Ar/Tt.Ar differences between immobilized and control limbs ranged from -7.4 to +0.6%, with the greatest decrease in C57BL/6J mice (Figure S1). C57BL/6J was the strain with the greatest difference between immobilized and control limbs for Ct.Ar/Tt.Ar, Ct.Ar, and Ct.Th.

**Figure 1.**
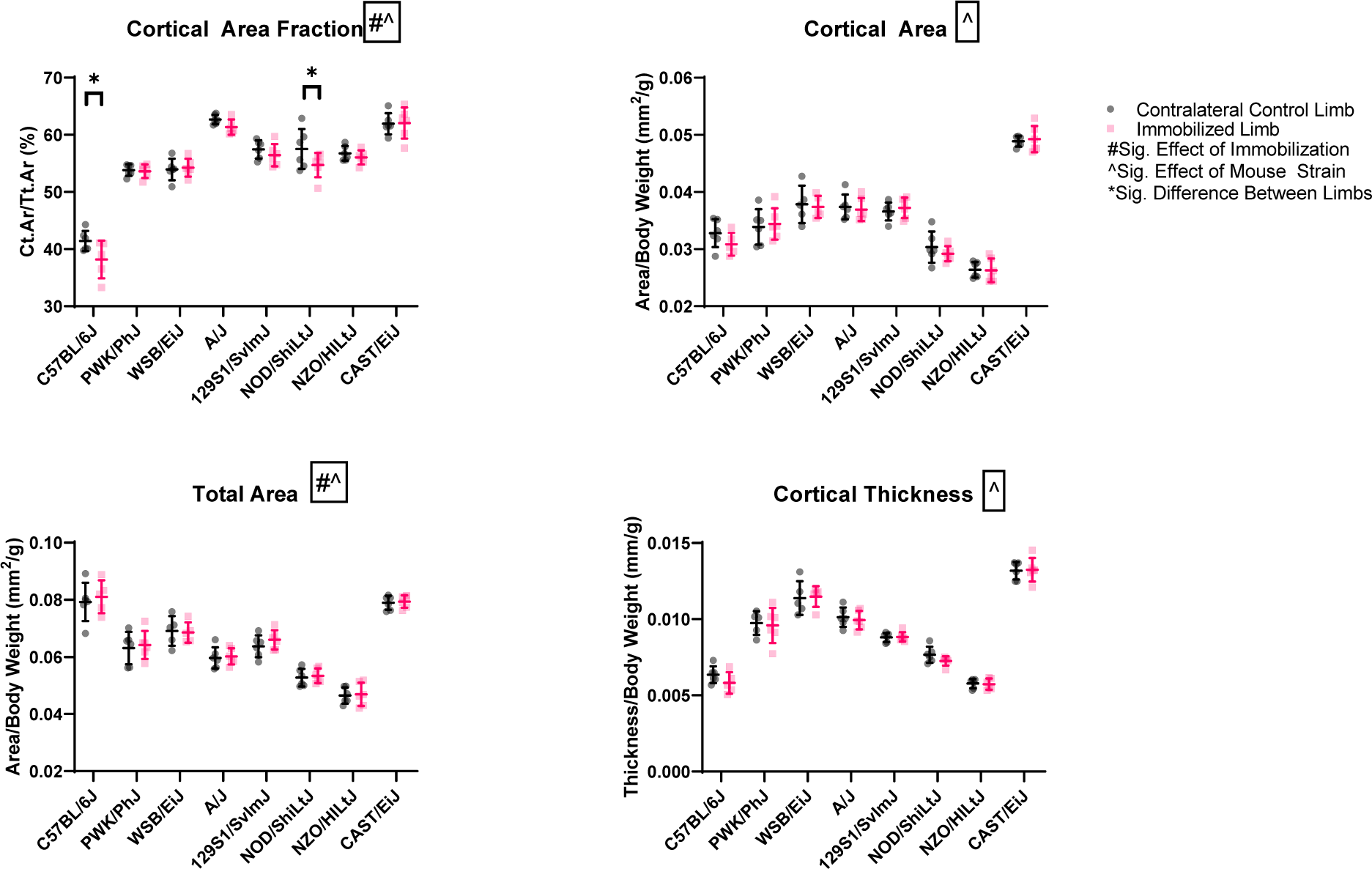
Femoral (mean ± SD) cortical bone of the mid-diaphysis after three weeks of single limb immobilization. Cortical area, total area, and thickness are normalized by body weight. There was a main effect of mouse strain on every property (p<0.0001), and there was a main effect of immobilization (p<0.05) on Ct.Ar/Tt.Ar and total area. C57BL/6J and NOD/ShiLtJ mice had decreased Ct.Ar/Tt.Ar from immobilization (p<0.05). C57BL/6J mice had the greatest magnitude of Ct.Ar/TAr loss. The other strains had no significant differences between immobilized and contralateral control limbs.

There were main effects of mouse strain on femoral metaphyseal BV/TV and Tb.N (p<0.0001, Figure 2). Immobilization resulted in a wide range of changes in BV/TV with some mouse strains losing bone and others gaining bone. NZO/HiLtJ mice had the greatest decrease of BV/TV with -11.9%, while A/J mice had the greatest increase with +16.1% (Figure S2). These two strains also had the greatest and lowest magnitude of effects for Tb.Th, Tb.N, and Tb.Sp.

**Figure 2.**
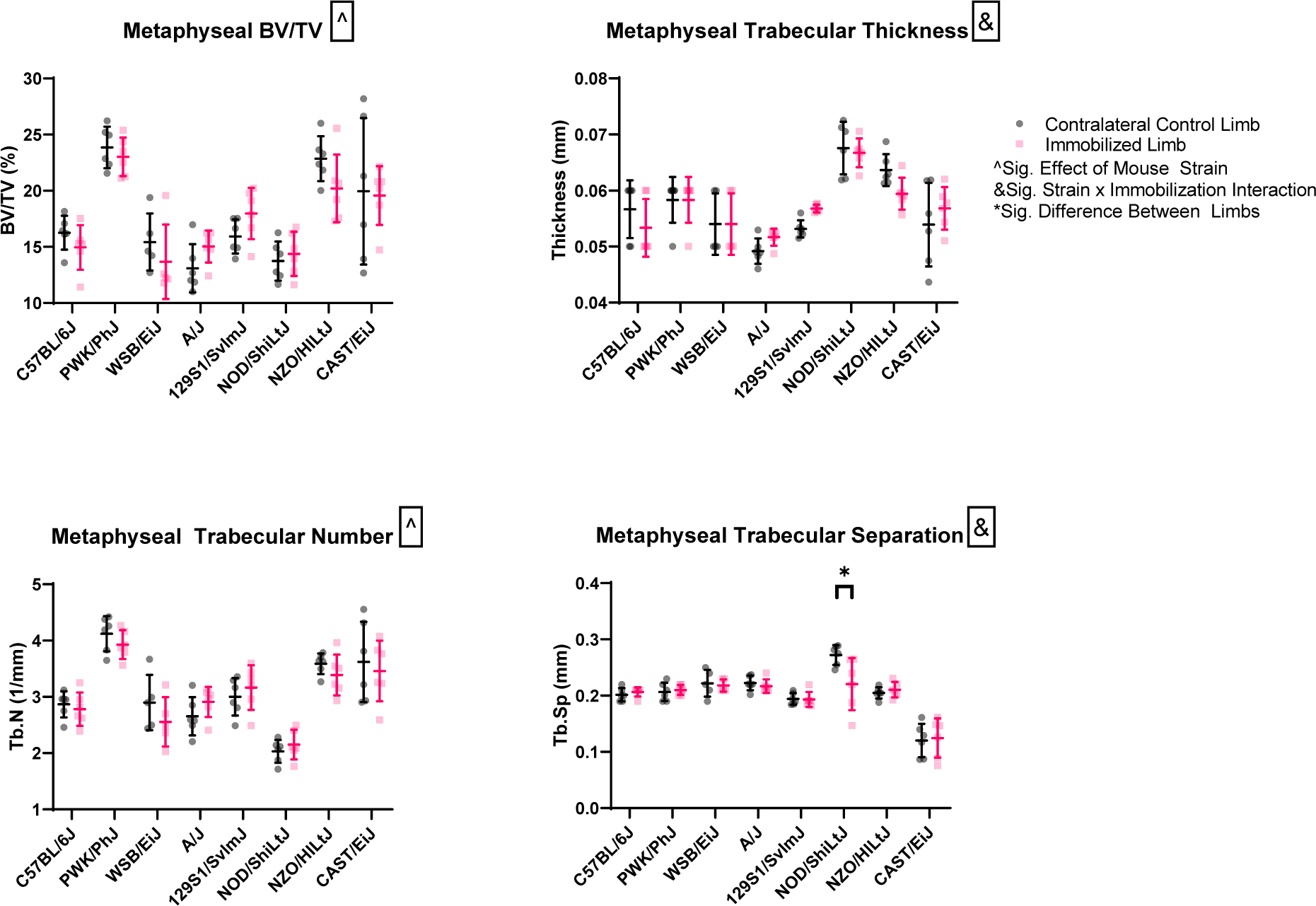
Femoral (mean ± SD) trabecular bone of the distal metaphysis after three weeks of single limb immobilization. There were main effects of mouse strain (p<0.05) on BV/TV and Tb.N. There was an immobilization and mouse strain interaction (p<0.05) for Tb.Th and Tb.Sp. All mouse strains had bone loss (p<0.05) in this region except A/J and 129S1/SvImJ.

In the femoral epiphysis, there were main effects of mouse strain and immobilization on Tb.Th and Tb.Sp (p<0.0001, Figure 3). Immobilization and mouse strain had interactive effects (p<0.0001) on BV/TV and Tb.N. Among the mouse strains, differences between BV/TV of immobilized and control limbs ranged from -36.5 to -8.3%, with the greatest decrease in CAST/EiJ mice (Figure S3). CAST/EiJ mice also had the greatest loss of Tb.N, while C57BL/6J mice had the greatest loss of Tb.Th. WSB/EiJ mice had the greatest increase in Tb.Sp.

**Figure 3.**
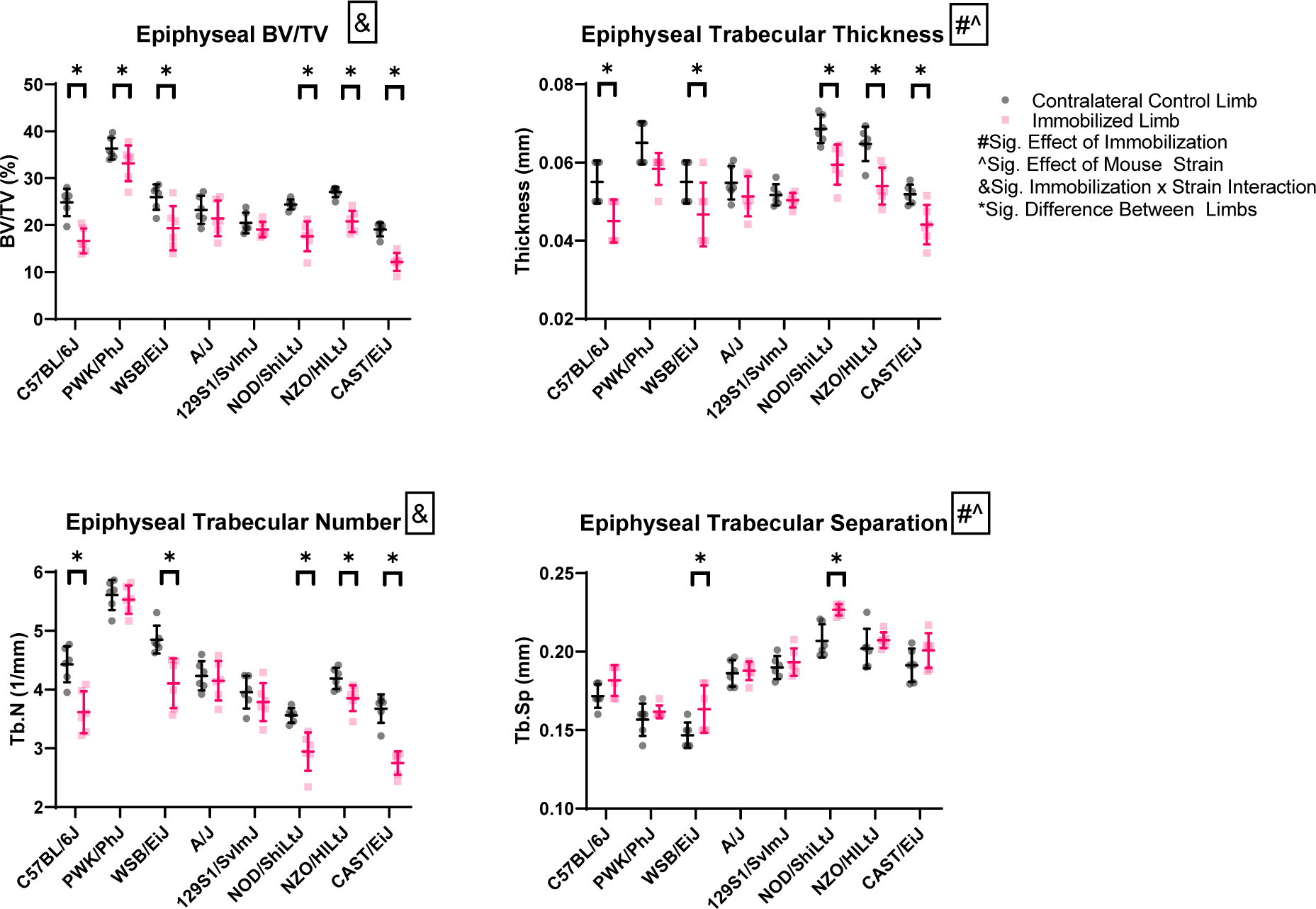
Femoral (mean ± SD) trabecular bone of the distal epiphysis after three weeks of single limb immobilization. There were main effects of immobilization and mouse strain (p<0.05) on Tb.Th and Tb.Sp. There was an immobilization and mouse strain interaction (p<0.05) for BV/TV and Tb.N. All mouse strains had bone loss (p<0.05) in this region except A/J and 129S1/SvImJ.

### Mouse Strain and Immobilization Influenced Bone Strength but to a Lesser Extent than Bone Geometry

There was a main effect of mouse strain on all femoral structural-level and tissue-level mechanical properties (p<0.0001, Figure 4). There was a main effect of immobilization on ultimate stress and elastic modulus (p<0.01). Effects of immobilization on mechanical properties were of much lower magnitude than bone geometry properties. Among the mouse strains, differences between ultimate stress of immobilized and control limbs ranged from -0.1 to +0.2% (Figure S4). C57BL/6J mice had the greatest loss of ultimate force, stiffness, ultimate stress and elastic modulus from immobilization.

**Figure 4.**
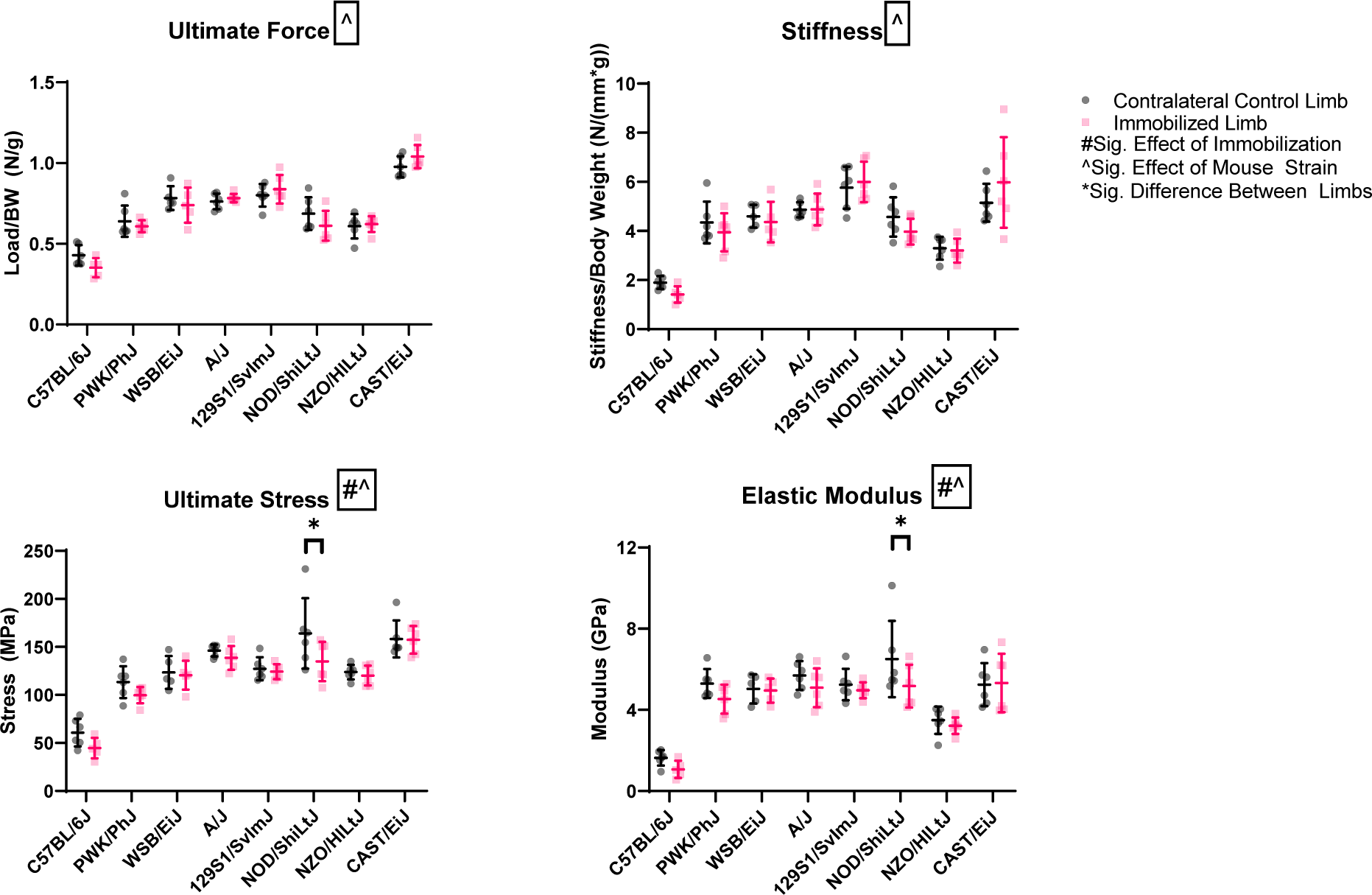
Femoral (mean ± SD) bone structural-level and tissue-level strength after three weeks of single limb immobilization, measured by three-point bending to failure. There was a main effect of immobilization (p<0.05) on ultimate stress and elastic modulus. There was a main effect of mouse strain (p<0.05) on all properties. Only NOD/ShiLtJ mice had loss of tissue strength (p<0.05) from immobilization.

### Mouse Strain and Immobilization Both Influenced Bone Chemical Composition

There was a main effect of mouse strain on Raman spectroscopy measures of mineral:matrix (p<0.0001), carbonate:phosphate (p<0.01) ratios, and crystallinity (p<0.0001, Figure 5). There was also a main effect of immobilization on these three outcome measures (p<0.0001). Among the eight mouse strains, differences between mineral:matrix of immobilized and control limbs ranged from -19.5 to -1.2%, with the greatest decrease in A/J mice (Figure S5). Immobilized limbs of 129S1/SvImJ, A/J, CAST/EiJ, NOD/ShiLtJ, PWK/PhJ, WSB/EiJ and NZO/HiLtJ mice had decreased mineral:matrix ratio. Immobilized limbs of A/J, C57BL/6J, 129S1/SvImJ, PWK/PhJ, NOD/ShiLtJ and CAST/EiJ mice had increased carbonate:phosphate ratio compared to control limbs. Crystillanity was the greatest in the control limbs of the A/J mice amongst the mouse strains, while C57BL/6J mice had the lowest. Crystallinity increased with immobilization in NZO/HiLtJ, PWK/PhJ, C57BL/6J, CAST/EiJ, 129S1/SvImJ and WSB/EiJ mice, but decreased in A/J and NOD/ShiLtJ mice.

**Figure 5.**
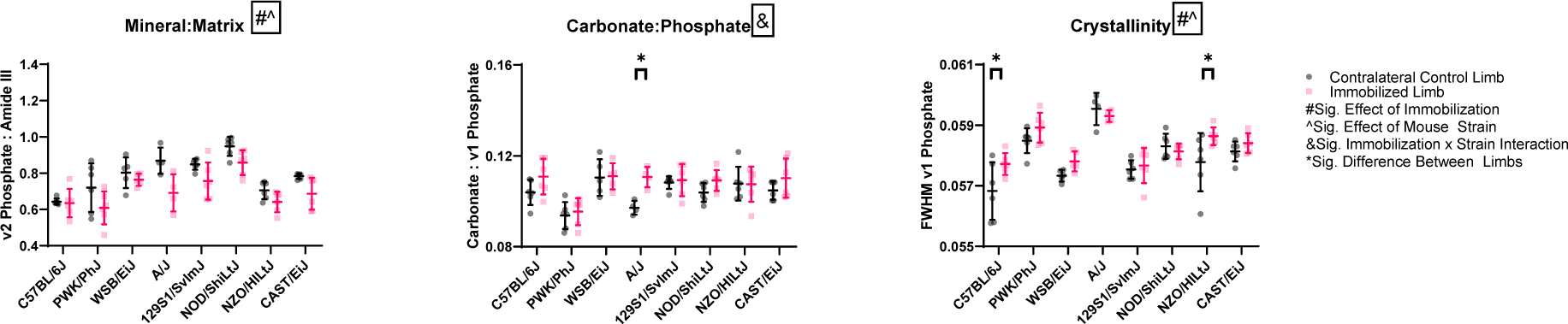
Femoral (mean ± SD) cortical bone mineral:matrix, carbonate:phosphate and crystallinity measured by Raman spectroscopy after three weeks of single limb immobilization. There was a main effect of immobilization (p<0.05) on mineral:matrix and crystallinity. There was a main effect of mouse strain (p<0.05) on all properties.

**Figure 6.**
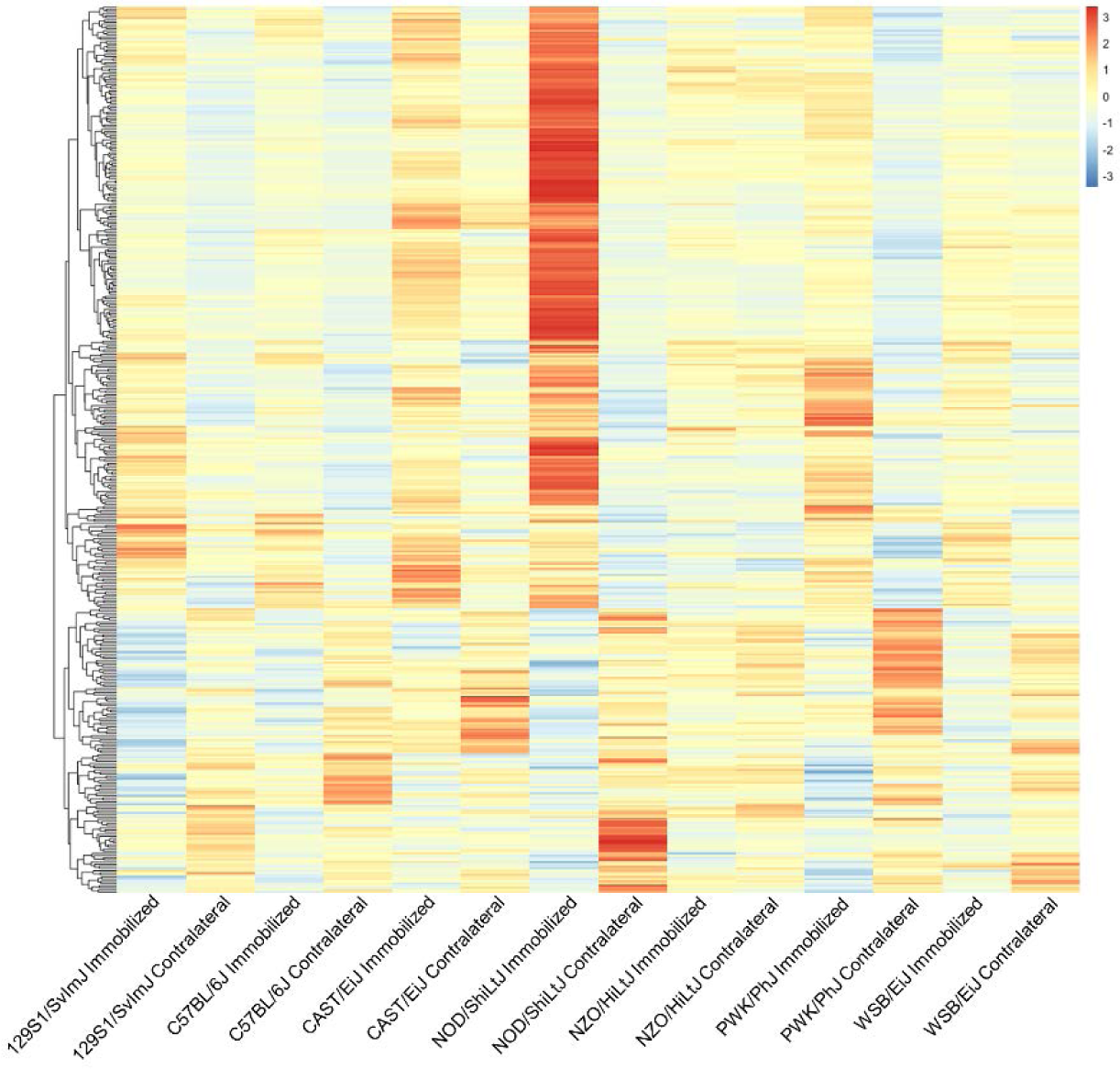
Heatmap showing relative gene expression of 414 genes affected by immobilization, as measured by RNA-seq ot tibial diaphyseal samples from immobilized and contralateral control limbs. Each row represents one gene. NOD/ShiLtJ and CAST/EiJ mice had the greatest number of genes affected by immobilization.

### RNA-seq Revealed Gene Expression was affected by Immobilization in Some but not All Mouse Strains after Three Weeks

There were 414 genes differentially expressed in response to immobilization across seven of the eight DO founders (Figure 7). CAST/EiJ mice had the greatest number of genes, 1095, differentially expressed between immobilized and control limbs. NOD/ShiLtJ mice had 788, and PWK/PhJ had 19 genes differentially expressed between immobilized and control limbs. No other strains had genes differentially expressed between immobilized and control limbs. Amongst the genes affected by immobilization, the genes with the highest expression in immobilized limbs were *Col1a2, Col1a1, Sparc, Spp1*, and *Col5a2*. Gene ontology analysis revealed the most highly upregulated pathways in immobilized limbs to be pathways involved in collagen formation, collagen assembly, and extracellular matrix organization (Table 2). There were 8,568 genes differentially expressed across the seven DO founders.

**Table 1.** Heritability of the response of each property to immobilization.

| Property | Heritability |
| --- | --- |
| Weight Loss | 62% |
| Cage activity | 24% |
| Food consumption | 21% |
| Ct.Ar/Tt.Ar | 31% |
| Ct.Ar | 15% |
| Ct.Th | 21% |
| Metaphyseal BV/TV | 27% |
| Metaphyseal Tb.Th | 35% |
| Metaphyseal Tb.N | 23% |
| Epiphyseal BV/TV | 64% |
| Epiphyseal Tb.Th | 35% |
| Epiphyseal Tb.N | 71% |
| Ultimate Force | 26% |
| Ultimate Stress | 28% |
| Mineral:Matrix | 16% |
| Carbonate:Phosphate | 30% |
| Crystallinity | 28% |

**Table 2.** Gene ontology analysis of the most highly upregulated pathways in immobilized limbs compared to contralateral control limbs.

| Reactome Pathways | (raw P-value) | (FDR) |
| --- | --- | --- |
| Collagen formation (R-MMU-1474290) | 2.01E-21 | 3.25E-18 |
| Collagen biosynthesis and modifying enzymes (R-MMU-1650814) | 2.54E-19 | 2.05E-16 |
| Extracellular matrix organization (R-MMU-1474244) | 2.02E-18 | 1.09E-15 |
| Assembly of collagen fibrils and other multimeric structures (R-MMU-2022090) | 1.08E-12 | 4.37E-10 |
| Degradation of the extracellular matrix (R-MMU-1474228) | 6.55E-11 | 2.12E-08 |

**Figure 7.**
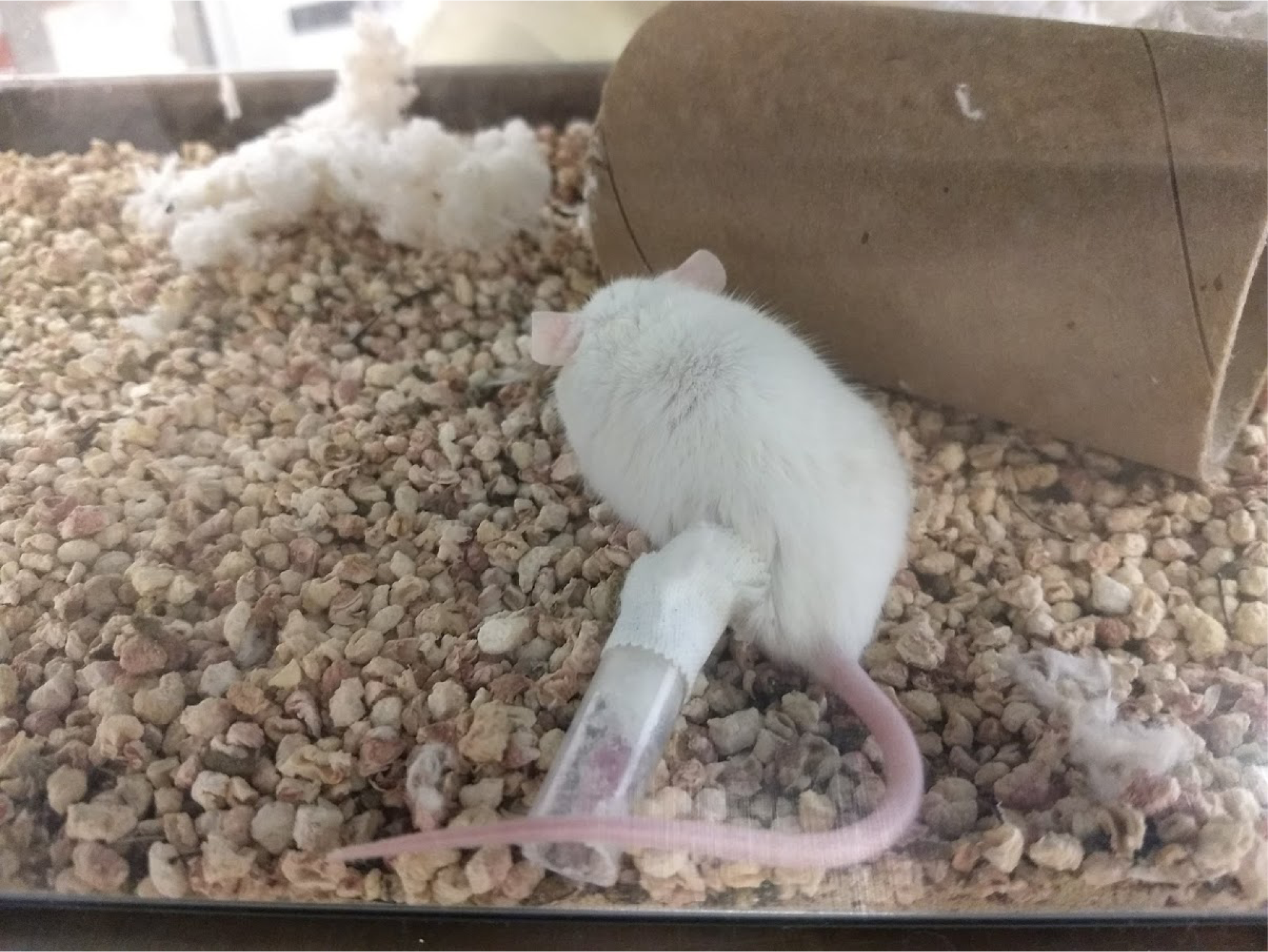
A/J mouse with its left hindlimb immobilized.

**Figure 8.**
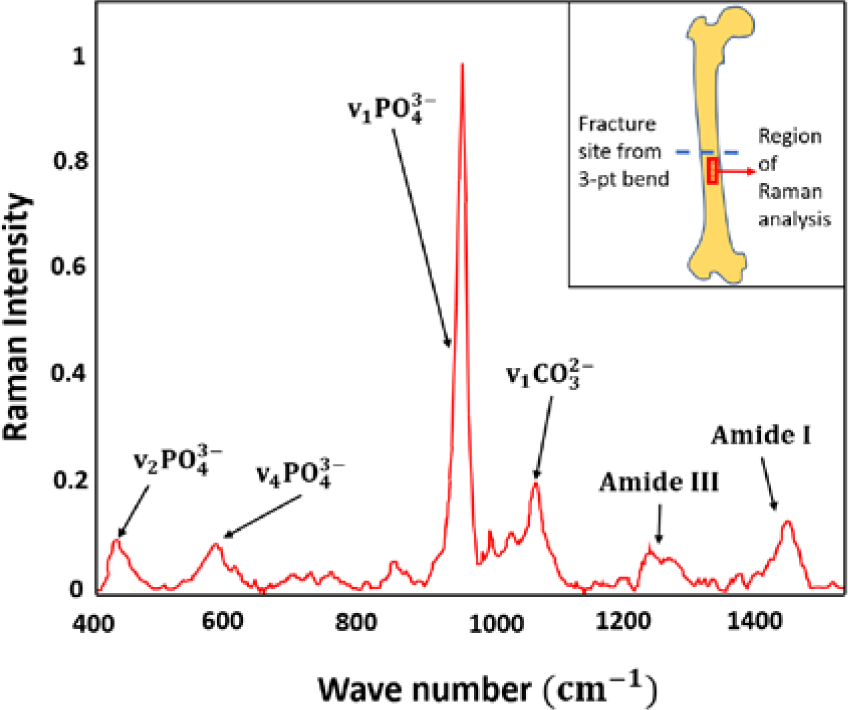
Example Raman spectrum of a mouse femur and position of the Raman array on the periosteal surface of the femur.

### All Strains Lost Body Weight from Immobilization

All strains lost body weight as a result of the immobilization (Table 3). Weight loss ranged from -2.6% body weight in PWK/PhJ mice to -17% in A/J mice. Mouse cage activity, measured in total distance traveled, decreased greatly in A/J mice after immobilization. All other strains had slight decreases or large increases in distance travelled. Food consumption increased in all strains except NZO/H1LtJ.

**Table 3.**
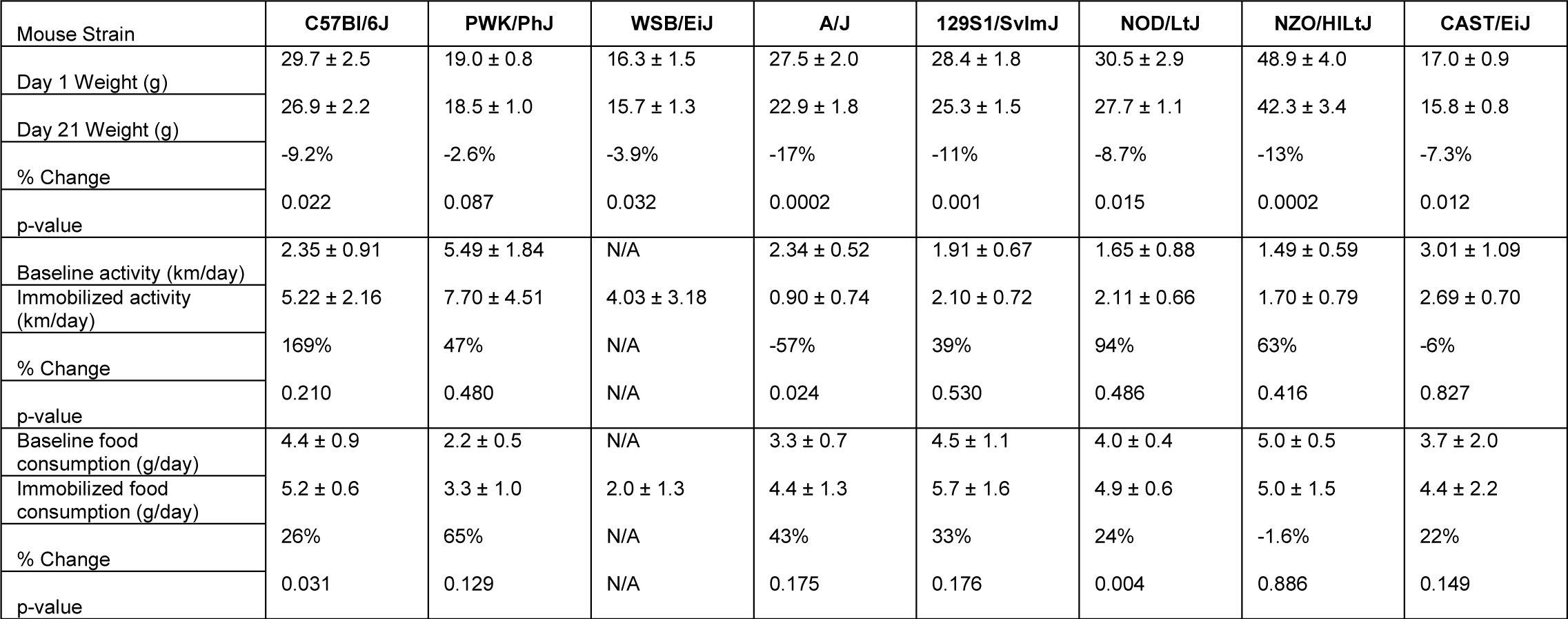
Mouse body weight, food consumption, and cage activity measurements.

### Heritability of Effects of Immobilization Varied by Region of the Bone

Heritability, the variance due to genetic factors, was calculated for each property measured as previously described (Table 1).^21^ Each heritability value shows the effects of genetic variation on the response to immobilization for that property. Loss of epiphyseal Tb.N had the highest heritability at 71%, followed by loss of epiphyseal BV/TV at 64%. This suggests changes in these properties in response to immobilization are influenced by genetic variation. Mineral:matrix had the lowest heritability at 16%, suggesting changes in tissue-level bone material properties in response to immobilization might not be as largely affected by genetic variation.

## Discussion

The DO mouse population is becoming a critical tool for evaluating the role of genetic variation on bone adaptation. This is the first study to examine the effects of mechanical unloding on the eight inbred DO founder strains. The greater number of strains used here than in previous unloading studies gives greater power to detect heritability.^22^ In this study, we observed that genetic background played a major role in the response of bone to immobilization. Some mouse strains were greatly affected while others demonstrated almost no negative effects of immobilization. This variety of effects suggests there may be some genes that protect bone from the negative effects of unloading. The DO mice that are derived from the founder strains studied here would also be expected to have a variety of responses to unloading, making them a suitable model for studying the effects of genetic variation. Heritability values for the responses of each property to immobilization ranged from 15-71%. This highlights the large systemic effects of genetic variation on the ability of bone to sense and respond to mechanical unloading.

C57BL/6J and NOD/ShiLtJ mice had the greatest diaphyseal cortical bone loss from unloading. As would be expected from these changes, these mice also had the greatest magnitude of loss of bone strength from unloading. NZO/HlLtJ mice had the greatest metaphyseal trabecular bone loss, and C57BL/6J, WSB/EiJ, NOD/ShiLtJ, and CAST/EiJ mice had the greatest epiphyseal trabecular bone loss. Regional differences in bone loss by strain may be due to differences in bone morphology, animal size, changes in gait, and activity levels, all of which are affected by genetic differences. That region-specific response to unloading may be influenced by genetics is an intriguing concept that requires further investigation.

The greatest overall effects of immobilization were seen in the femoral epiphysis as every strain had loss of BV/TV in this region. Epiphyseal bone may be uniquely positioned to suffer bone loss from immobilization as this region has a greater amount of trabecular bone, and epiphyseal bone could experience a large decrease in mechanical loading following the removal (or reduction) of joint contact forces. Unloading also reduces chondrocyte growth and differentiation which may negatively affect bone metabolism in the epiphysis.^23,24^ Bone loss in the epiphysis had the highest heritability, suggesting a greater influence of genetics in this region. There is sparse data on epiphyseal bone changes in response to unloading in the literature.^25^ As several factors may affect bone metabolism in the epiphysis, the epiphysis may be a region of high interest for future research on effects of unloading on bone.

While bone quality measures have been linked to genetic background in mice,^26^ this is the first study to examine bone quality measurements in a large number of genetically diverse, inbred mouse strains exposed to mechanical unloading. Mineral chemistry measures varied greatly across the eight strains, and bone quality responded to immobilization in a strain-specific manner. With immobilization, bone chemistry measures followed a general pattern of lower mineral:matrix, increased carbonation (carbonate:phosphate), and increased crystallinity. Unloading at osteoporotic fracture sites in postmenopausal women resulted in diminished mineral:matrix.^27,28^ Similarly, growing rats exposed to 12.5 days of microgravity on COSMOS 1179, a spaceflight experiment, had reduced apatite mineral and bone stiffness.^29^ Such compositional changes in bone, along with increased carbonate:phosphate and crystallinity with immobilization, may contribute to decreased bone strength. Carbonation, which increases with tissue age, has been shown to increase in femoral trabecular bone of women who had an osteoporotic fracture and in adult female rats following hindlimb unloading.^27,30^ Increased substitution of carbonate in the hydroxyapatite crystals in the bone can cause unsymmetry in the crystal lattice, making the bone more prone to damage under mechanical stress.^31,32^ Smaller mineral crystals enable greater deformation of mineralized collagen fibers, thus leading to higher post-yield deformation and reducing propensity toward fracture. In contrast, increased crystallinity contributes to poor bone strength where larger crystals increase the brittleness of bone.^33^ Crystallinity and brittleness has also been shown to increase with age^34^ and, in some studies, with unloading from microgravity exposure.^29,30,35^ Thus, these changes in bone chemistry may contribute to impaired fracture toughnening mechanisms and reduce bone strength. As the changes to bone chemistry and bone geometry measures due to immobilization are strain-specific,^26^ this indicates that the DO population holds potential to reveal specific genes that may play roles in preventing detrimental bone chemistry alterations that occur with disuse.

Changes in gene expression in response to unloading were greatest in NOD/ShiLtJ and CAST/EiJ mice. These results are different from diaphyseal cortical bone geometry data that showed large effects of unloading in C57BL/6J mice. This may reflect a timing issue where C57BL/6J mice have more differentially expressed genes in response to unloading at an earlier time point and have reached some homeostasis after three weeks. Measuring gene expression data does have the limitation of only analyzing expression at one moment in time when gene expression fluctuates greatly throughout the day and from week to week. Additionally, we measured gene expression in the tibia and bone geometry in the femur, so there may have been bone-specific differences in the response to unloading. Alternatively, differences in measurement variance within each strain may have made significant differences difficult to detect for this study’s sample size. All of these differences highlight the influence of genetic variation on the timing and location of the skeletal response to unloading.

Interestingly, the gene ontology pathways shown to be most highly upregulated in response to unloading were osteogenic pathways associated with increasing collagen synthesis and extracellular matrix formation. This is similar to the response seen after anabolic mechanical loading.^36-38^ It is possible that at the time point analyzed, bones had reached a homeostasis where they were no longer perceiving a decrease in mechanical loading and were beginning to remodel. Although there was increased expression of osteogenic genes, the new bone formation in immobilized bones is likely to be of lower quality than in the controls.^39,40^ This was one of the first studies to use RNAseq to examine changes in gene expression in bone in response to unloading and to compare gene expression across multiple mouse strains.^41^ There were some limitations as we were only able to analyze seven of the mouse strains, and gene expression was analyzed in the tibia while all other measurements were done in the femur. Further studies need to be performed to better understand the time course of changes in gene activation with changes in mechanical loading and how genetic variation affects these responses.

We did not detect changes in physical activity in the mouse strains that had the greatest bone loss. Physical activity increased in most strains after immobilization. Although the casts made ambulation more difficult and cumbersome, the mice appeared to be more active by trying to free themselves from the casts. A/J mice had decreased activity, but that did not translate to bone loss as A/J mice were one of the least affected strains. This can be a limitation of the activity level measurement which only showed distance traveled and did not take into account changes in gait or speed which could affect bone loss.^42^ Surprisingly, food consumption increased for most of the strains, a result that would not be expected to be associated with bone loss.^43^ Other changes in mouse behavior not measured may be able to partially explain the bone loss from immobilization in this study.

A major limitation of this study was the use of inbred mice to evaluate the effects of genetic variation. Inbred mice have limited power to detect these effects. Future work should be performed using outbred mice. Since differences in the response to immobilization were found amongst the eight DO founder strains, we would expect to see differences in the response to immobilization in DO mice.

Genetic variation plays a major role in the accrual and maintenance of bone mass. Here we showed genetic variation in mice resulted in differences in the skeletal response to unloading from immobilization. There were mouse strain-dependent differences in gene expression and in changes to bone volume, bone strength, and bone chemical composition from immobilization. We can next evaluate the response to immobilization in DO mice and use that information to uncover novel genes involved for protecting bone from the negative effects of unloading.

## Materials and Methods

### Animals

All animal procedures were performed with the approval of the Virginia Commonwealth University Institutional Animal Care and Use Committee. Six male mice of each of the DO founder strains were purchased from the Jackson Laboratory (PWK/PhJ – stock #003715, WSB/EiJ – stock #001145, A/J – stock #000646, 129S1/SvImJ – stock #002448, NOD/ShiLtJ – stock #001976, NZO/HlLtJ – stock #002105, and CAST/EiJ – stock #000928) at between four and fourteen weeks of age. Mouse age, sex, sample size, and immobilization duration were determined based on previous results using single limb immobilization.^20^ C57BL/6J (stock #000664) bone properties and bone samples collected from that study were also included in the analysis for this study. All mice were kept in single housing upon arrival and for the duration of the study. At sixteen weeks old, all mice had casts placed on their left hind limbs. Mice were sacrificed after three weeks in the casts.

### Casting Protocol

The left hind limb of each mouse was immobilized in a cast as previously described.^20^ Mice were placed under general anesthesia, and surgical tape was wrapped around the left hind limb. A microcentrifuge tube, with the bottom end removed, was then glued onto the tape. The contralateral right hind limbs were not altered and were used as controls.^18^ All mice were able to move around the cages by dragging the immobilized limb or rotating the limb at the hip (Figure 7).

### Physical Activity and Food Consumption

Physical activity and food consumed was observed over 24-hour periods, before and after immobilization. These observations were performed one week prior to casting and two weeks after casting. Video was recorded for 24 hours and analyzed using the OpenField Matlab function developed by Patel *et al*.^*44*^ Total distance traveled over 24 hours was measured for each mouse. Food consumption was measured by weighing food in the cages before and after the 24-hour observation period.

### Bone Morphology

Femurs were harvested at sacrifice and stored at -20°C in calcium buffer. The bones were scanned by micro-CT (Skyscan 1173, Bruker microCT) as previously described.^20^ Bones were scanned at 8.6 µm voxel size, 70kVp, 114 microA, 1.0 mm Al filter, and 1200 ms integration time. A 180-µm slice of the mid-diaphysis was used to measure cortical area (Ct.Ar), area fraction (Ct.Ar/Tt.Ar), thickness (Ct.Th), and tissue mineral density (TMD). Metaphyseal trabecular bone morphology was evaluated using a 750-µm slice taken 200 µm proximal from the growth plate. Computer-traced ROIs were used to measure bone volume fraction (BV/TV), trabecular thickness (Tb.Th), trabecular number (Tb.N), trabecular separation (Tb.Sp), and TMD.^45^ Epiphyseal trabecular bone morphology was measured using a 520-µm slice of the epiphysis with freehand-traced ROIs immediately distal to the growth plate.

### Mechanical Properties

All femurs were tested by three-point bending to failure under displacement control at 1.0 mm/min. Bones were loaded with the anterior side in tension on support spans of 8 mm. Stiffness (instantaneous slope of the linear portion of the load-displacement curve at 3.5 N load), yield load (load at loss of 10% of stiffness), ultimate load, yield displacement, ultimate displacement, and work were measured.^46^ Geometry measurements at the location of the fracture site in the micro-CT scans were used to normalize structural-level properties to give estimates of elastic modulus, yield stress, ultimate stress, yield strain, ultimate strain, and toughness.^47^

### Raman Spectroscopy

Raman spectroscopy was performed on the periosteum^48^ of the distal aspect of the mid-diaphysis of the femurs, on the anterior side, 0.5 mm from the fractured surface from 3-point bending, with a Renishaw inVIA confocal Raman spectroscope (Renishaw, Wotton-under-Edge, Gluocestershire, UK; 785 nm wavelength laser, 6 s exposure time, 10 accumulations and 100% intensity). Bones were wrapped in gauze soaked in calcium buffer and were stored at -20°C. Prior to data collection, samples were thawed for 1 hour and gently cleaned with a soft toothbrush to remove remaining soft tissue. The samples were not otherwise polished. Raman measurements were collected in a 2D map (3 × 20 sites with 20 μm spacing) that was aligned along the bone’s long axis.^49^ All samples were evaluated while submerged in PBS. Peaks were centered around 1000 Raman shift/cm. The fluorescence baseline from all the spectra were subtracted, using an 11th order polynomial fit to the entire spectrum, followed by cosmic ray removal in the Renishaw Wire software. These spectra also underwent normalization. Custom MATLAB code was used to calculate peak area ratios of v2 phosphate:amide III or mineral:matrix ((413-466)/(1218-1309)cm^-1^), carbonate:v1 phosphate or carbonate:phosphate ((1057-1104)/(917-984)cm^-1^), and crystallinity (inverse of half-width at full maximum height of the v1 phosphate peak at 959cm^-1^ as in Figure 9.^50-52^ Data points from samples with low signal-to-noise ratios or data points which fell on residual soft tissue were removed and were excluded from the analysis.

### RNA Sequencing

The tibial diaphysis, without marrow, from immobilized and contralateral mice from the 8 DO founders (n = 3 per strain/group) was isolated and total RNA isolated using Trizol. RNA quality was assessed using an Agilent 4200 TapeStation System. RNA was of sufficient quality (RIN>5) for all samples except from the A/J strain. RNA-seq libraries were generated using Illumina TruSeq Stranded mRNA Library Prep kits and sequenced on an Illumina NextSeq500 for the seven strains with high-quality RNA. Raw reads were aligned to the reference Mus Musculus genome (GRCM38) using Hisat2.^53^ Transcript assembly and quantification was conducted using StringTie.^54^ Differential Gene Expression analysis was completed using EdgeR.^55^ Gene Ontology (GO) terms were assigned using PANTHER (Fisher test with FDR correction at 0.05).^56^

### Statistical Analysis

Repeated measures two-way ANOVAs were used to test for significant (p<0.05) main effects of mouse strain, immobilization, and interactions. Sidak tests post-hoc were used to test for significant differences between control and immobilized limbs of each strain. For Raman analysis, as multiple test sites were collected on each bone from Raman spectroscopy, a repeated measures mixed model analysis was performed in software JMP 14, with mouse strain (8 levels) and loading condition (2 levels) as fixed effects (with interaction term) and mouse number as a random effect. Tukey’s post hoc was used to test for significant differences in various Raman parameters for the two fixed effects. Cortical bone geometry and structural bone mechanical properties were normalized by body weight before testing for effects of mouse strain.^46^ Magnitude of effects of immobilization were determined by calculating percent differences between immobilized and contralateral control limbs. One-way ANOVAs with Tukey’s tests post-hoc were used to test for significant differences in magnitude of effects of immobilization. Heritability, the variance due to genetic factors, was calculated for the magnitude of effects of immobilization on each property measured as previously described.^21^ The R-squared values from the One-way ANOVAs were used as the heritability values since they represent the variation in the data from mouse strain.

## Acknowledgments

This work is supported by the National Institutes of Health (R01AR068132-20, 5R01AR068345, R01AR071657, T32LM012416), National Science Foundation (NSF CBET 1338154), National Aeronautics and Space Administration (80NSSC18K1473), and the Translational Research Institute for Space Health Postdoctoral Fellowship (NASA Cooperative Agreement NNX16AO69A).

## Conflicts of Interest

None.

## Contributions

MAF, VLF, CRF, and HJD contributed to study design. MAF, AA, BS, YZ, and CRM contributed to data acquisition. MAF, AA, BS, VLF, CRF, and HJD contributed to interpretation of results. MAF, AA, BS, VLF, CRF, and HJD contributed to drafting and revising the manuscript.

## Notes

### Competing Interest Statement

The authors have declared no competing interest.

